# A revised single-cell transcriptomic atlas of *Xenopus* embryo reveals new differentiation dynamics

**DOI:** 10.1101/2024.01.02.573882

**Authors:** Kseniya Petrova, Maksym Tretiakov, Aleksandr Kotov, Anne H. Monsoro-Burq, Leonid Peshkin

## Abstract

This paper introduces an updated single-cell atlas for pivotal developmental stages in *Xenopus*, encompassing gastrulation, neurulation, and early tailbud. Notably surpassing its predecessor, the new atlas enhances gene mapping, read counts, and gene/cell type nomenclature. Leveraging the latest Xenopus tropicalis genome version, alongside advanced alignment pipelines and machine learning for cell type assignment, this release maintains consistency with previous cell type annotations while rectifying nomenclature issues. Employing an unbiased approach for cell type assignment proves especially apt for embryonic contexts, given the considerable number of non-terminally differentiated cell types. An alternative cell type attribution here adopts a fuzzy, non-deterministic stance, capturing the transient nature of early embryo progenitor cells by presenting an ensemble of types in superposition. The value of the new resource is emphasized through numerous examples, with a focus on previously unexplored germ cell populations where we uncover novel transcription onset features. Offering interactive exploration via a user-friendly web portal and facilitating complete data downloads, this atlas serves as a comprehensive and accessible reference.

## Introduction

The ability to characterize gene expression at the single-cell resolution has revolutionized our understanding of embryogenesis and cell type differentiation. Recently, a group of leading specialists formulated guiding principles for single-cell-centered medical research as part of the LifeTime Initiative [1]. This initiative aims to track, understand, and target human cells during the onset and progression of complex diseases, analyzing responses to therapy at single-cell resolution. Applying this strategy to key medical challenges is anticipated to pave the way for cell-based interceptive medicine in the next decade.

Observing biological systems at the cellular level offers an unprecedented opportunity to define the functional modularity and combinatorial interactions of genes in various physiological contexts. Many of these contexts are evolutionarily conserved, establishing a baseline from which deviations can lead to malformations and disease. Currently, a Human Cell Atlas [2] is under construction, intended to form the core of this new single-cell perspective. However, research cannot solely rely on patients or human cell lines; biomedical science critically depends on model organisms.

Gene functions in the context of cell types are elucidated through cross-organism comparisons. Parallel work in model organisms, such as mice [3], adult flies, and zebrafish embryos [4,5], involves constructing cell atlases. In addition to immediate data analysis, these atlases serve as crucial reference resources and baselines for ongoing work. The value of single-cell transcriptomics resources in non-model organisms depends critically on the quality of the genome sequence and its annotation.

Since publishing the original atlas of embryonic development in Xenopus [6], significant improvements, including a chromosome-level assembly and gene annotation of the Xenopus genome, additional sequenced libraries, and more efficient pipelines for read mapping and expression analysis, have been released. This paper presents an updated and improved version of the Xenopus embryonic cell atlas, referred to as “XA23” for Xenopus embryonic Atlas 2023. Since the previous edition of the atlas is widely used, we begin with a re-analysis of previously published cell type nomenclature and translate it into the new version based on the latest genome annotation iteration, comparing it with findings from the current genome and annotation methods, with a specific focus on a novel population of germ cells.

## Results

### The new Embryonic Atlas supersedes the last version in per-stage and per-cell type coverage

For the new version of the *Xenopus* tropicalis single-cell atlas, we re-processed the inDrops (v2, v3) libraries from Briggs et al. [6] (dubbed “XA18”) and re-sequenced additional inDrops v3 libraries. The reads were aligned to the most recent version of *Xenopus* tropicalis genome (v10.7). As a result, the new atlas contains both the cells which were already present in the previous version (we call “old cells”) identifiable by unique barcodes, and the additional cells (we call “new cells”). There were numerous reasons for the new cells to appear. First, some cells were obtained from the resequenced data representing new library material. Second, the improved version of the genome allowed the assignment of more reads to the protein-coding genes not represented in the previous version of the genome. Finally, some of the cells with relatively low total gene expression were added since we identified these as legitimately low-expressing cells rather than cells which lost mRNA in the pipeline as we detail below.

Our data covers *X.tr.* embryonic development stages with even numbers from NF8 to NF22 [7] and additionally stages NF11 and NF13. After filtration, we recovered a total of 188,618 cells with a median of 1,269 UMI (Fig. 1a/b) and 600 genes per cell (Fig. S1) as compared to 136,966 cells, 1466 UMI and 764 genes per cell in the first atlas release XA18. Most stages received more cells, particularly benefiting representation of stages NF11 and NF8. In all stages some old cells did not pass the new quality filters, most notably in stage 13, but there is uniformly a net gain once we take the new cells into account. Of note, retained old cells have mostly gained in coverage, particularly non surprisingly in stages 11 and 13 - these stages were profiled only via inDrop v3 at low coverage in the original atlas. The new cells in all stages are of coverage lower than the rest of the cells (Fig. 1a), which is justified because these cells find reliable cell type attribution (Fig. 1c). In many cases we substantially improved the population size which should enable new insights into differentiation. Specifically, we have substantially expanded the representation of the lens placode, ionocytes, cardiac mesoderm and Spemann organizer (Fig. 1c) while reducing the representation of cranial neural crest, the hindbrain and trigeminal placodes.

**Figure 1.**
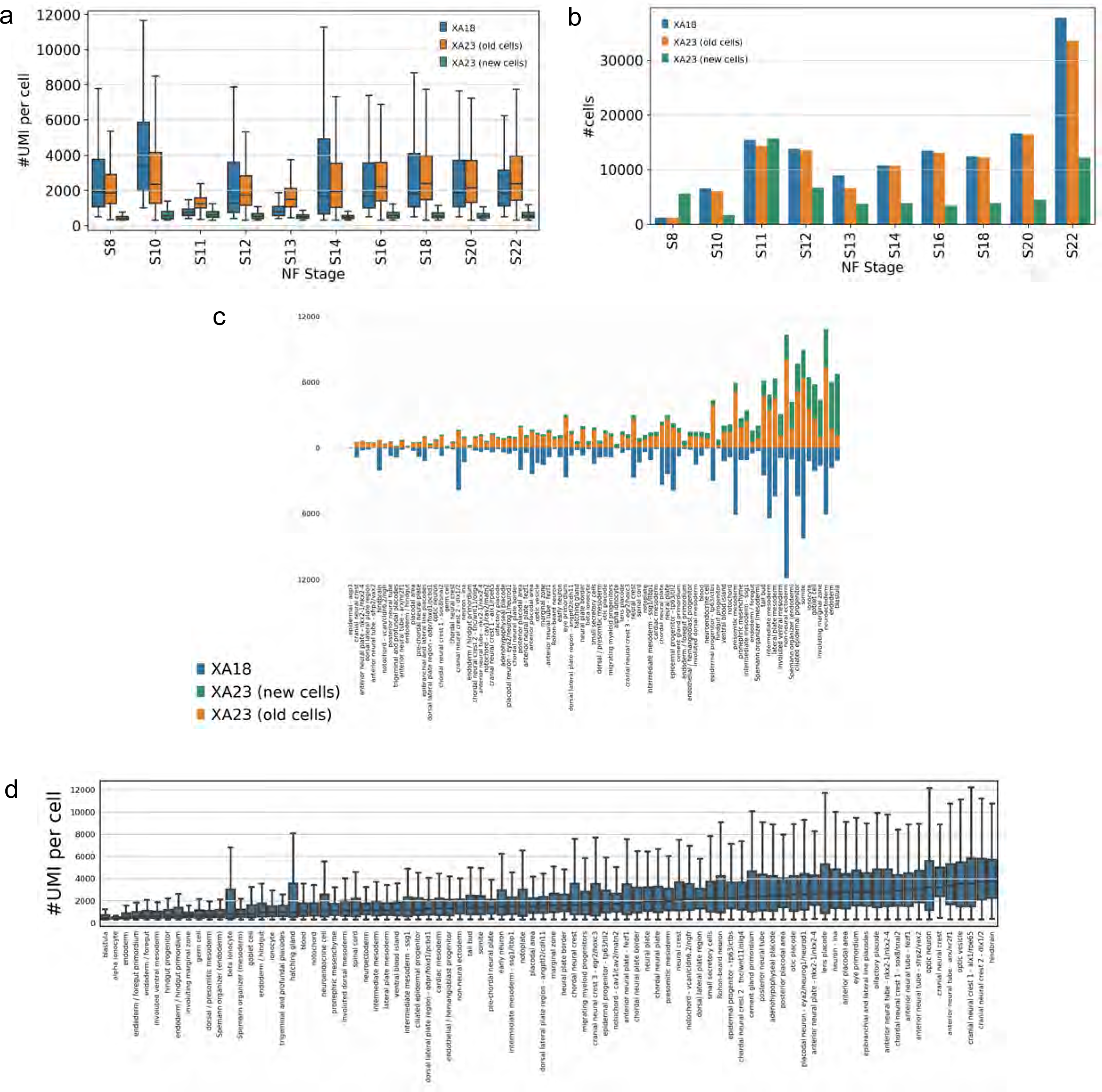
Comparison and summary statistics contrasting the XA18 and XA23 datasets. **(a)** The depth of coverage - number of UMI per cell. **(b)** The total number of cells per stage. **(c)** Cell-type specific coverage comparison. **(d)** The relative depth of coverage per cell type - germ cells have much less mRNA than neurons.

The depth of coverage is largely defined by specific processing pipeline and respective alignment parameters, yet some reduction in average UMI coverage should be attributed to inclusion of cells with lower coverage as justified here. Note that customary filtering of cells based on thresholding the total mRNA levels does not take into account the fact that cell types differ widely in how much mRNA they harbor. Cells that are actively synthesizing proteins will have a lot of mRNA [35,36]. This includes cells that are undergoing rapid growth and division, cells that are responding to stimuli or stress, and cells that are specialized for protein synthesis, such as secretory cells and neurons. The distribution of UMI per cell type, naturally reflects e.g. the mRNA abundance in neuronal cell types and low mRNA abundance in germ cells and endoderm types such as gut progenitors and specialized cells like ionocytes (Fig. 1d).

We assessed both whether the previous cell type annotation works well using the new genome annotation version and annotated the new cells, not presented in the XA18. Overall we kept most cell types but respective representation has changed.

### The new Embryonic Atlas is available for interactive Web browsing and interrogation

We made the realigned data available for browsing in the Spring [8] web browser, provided for each gene its *X.tr.* gene ID, name of corresponding gene in *X.tr.* v9.0 genome annotation and name of human best-hit homolog gene, if available, e.g. “ddx25 | Xetrov107048846m”. As previously described [6,9] the browser provides a wide range of functionalities for data interrogation, such as selecting subsets according to gene expression or pre-defined cell type, comparing sets of cells to find differentially expressed genes, creating and sharing derivative force-directed layouts, etc. For convenience of time-series expression analysis we grouped stages into three time adjacent phases with overlap: gastrula (St8-12), neurula (St 12-18) and tailbud (St 18-22). One new feature we introduce in this version is a secondary soft cell type index. I.e. in addition to assigning each cell a type label of deterministic nature, we calculate an index which assigns every cell to every cell type with some certainty. This is often a more natural situation in an embryo where only in rare cases there are terminally differentiated cells like e.g. cement gland or hatching glad, yet in most cases the cells are on a differentiation path where decisions have not been made e.g. cranial vs chordal neural crest. Selection of the cells for further analysis can be done using the adjustable range brackets over the value of cell type index, in order to focus on terminally differentiating or transient expression states.

*N.B. In the interest of keeping a focused discussion, most analysis below this point is focused on St 18 as a representative state, the other developmental states produce similar outcomes (data not shown)*.

### The original cell type annotation was largely sound yet somewhat deficient

We started with a consistency assessment of the XA18’s cell type annotation. We wanted to know whether cells which received a certain cell type label based on the gene marker analysis and XAO [10] constitute consistent annotation where cells of a given cell type are similar to one another and dis-similar to other cell types. We assessed the cell type annotation in the XA18 using two distinct approaches from classical Machine Learning. First, we built a kNN (nearest neighbor) cell type classifier, where we withheld the cell type label and compared it with what is predicted based on several cells most similar in expression to a given cell (see Methods). The consistency of the cell type annotation was thus verified by standard machine learning approach - cross validation, i.e attempting to predict a cell type of cells withheld from training the ML model. The prediction TPR (true positive rate) of labels averaged by cell type was 0.8±0.1 via 3-fold CV (cross-validation). As an alternative approach, we used a marker gene-based (dubbed mgB) prediction where the cell was assigned a cell type based on the expression of several key marker genes as defined in the original atlas annotation (see Methods). The classification TPR averaged by cell type was 0.6±0.2 via 3-fold CV (Fig. 2).

**Figure 2.**
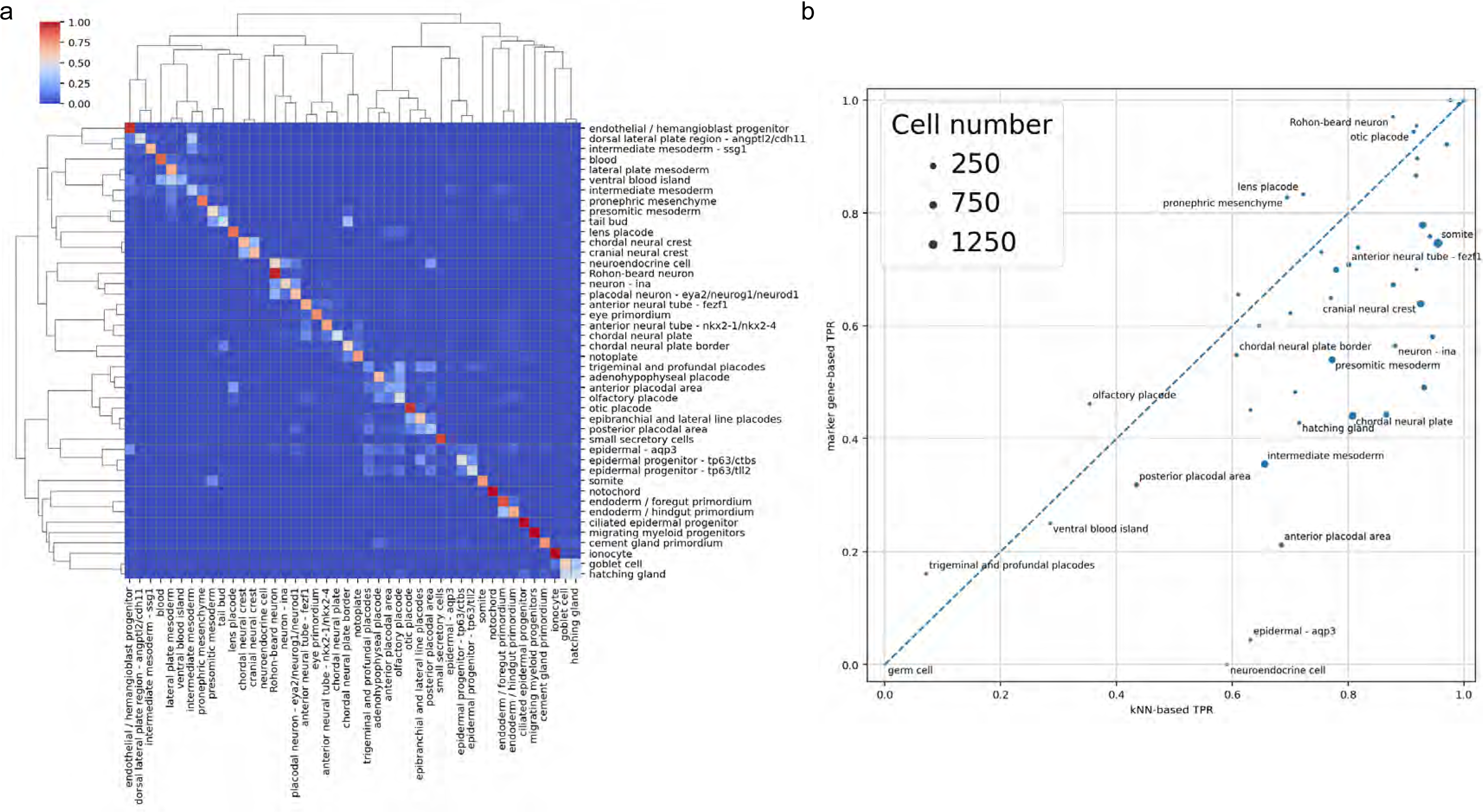
An assessment of cell type annotation consistency. **(a)** A confusion matrix for NF18 cell types prediction with marker gene-based approach in Briggs et al. dataset, annotated (rows) versus predicted (columns) cell types. **(b)** mgB prediction TPR vs. kNN prediction TPR for different NF18 cell types, disc sizes are proportional to cell type abundance.

We decided to explore this two-pronged approach to cell type annotation assessment because on one hand kNN results can be easily compared across the previous and the current genome annotations, on another the mgB approach is readily interpretable and allows us to validate the clusters in the sense of biological function and embryological context. As could be seen (Fig. 2a) while most cell type annotation turns out to be fairly consistent, there are some cell types which turned out to be poorly predictable with both kNN and gene markers. Not surprisingly, related cell types are intermixed, e.g. chordal and cranial neural crest, two subtypes (*tll2* and *ctbs*) of epidermal progenitors and foregut/hindgut primordium. Consider (the bottom left corner of Fig. 2b) e.g. germ cells, trigeminal and profundal placodes, ventral blood island and posterior placodal. We expected the germ cell type to be poorly predictable because, in the original atlas dataset, there were only 3 cells of this type. The rest of the poorly predictable cell types do not seem to have any strongly upregulated marker genes (Fig. 3b). Some cell types are predicted reasonably well by the kNN (TPR > 0.5) while being quite poorly predicted via marker genes (TPR < 0.2), e.g. neuroendocrine cells, epidermal-aqp3 and hatching gland (Fig. S2a). Only the “neuroendocrine cell” seems to have distinctive marker genes among these. Still, the limited set of upregulated genes and a small total number of cells (11) doesn’t allow us to reliably define these cells as neuroendocrine. The remaining cell types are relatively well predicted by both kNN and mgB (both TPR > 0.5) (Fig. 3a, Fig. S3). For example, the notochord is specifically and uniquely marked by the chordin (*chrd*), fibrillin (*fbn3*) and elastin microfibril interfacer (*emilin3*), while beta and gamma subunits of *Sec61* translocon complex are highly expressed but not specific to the notochord.

**Figure 3.**
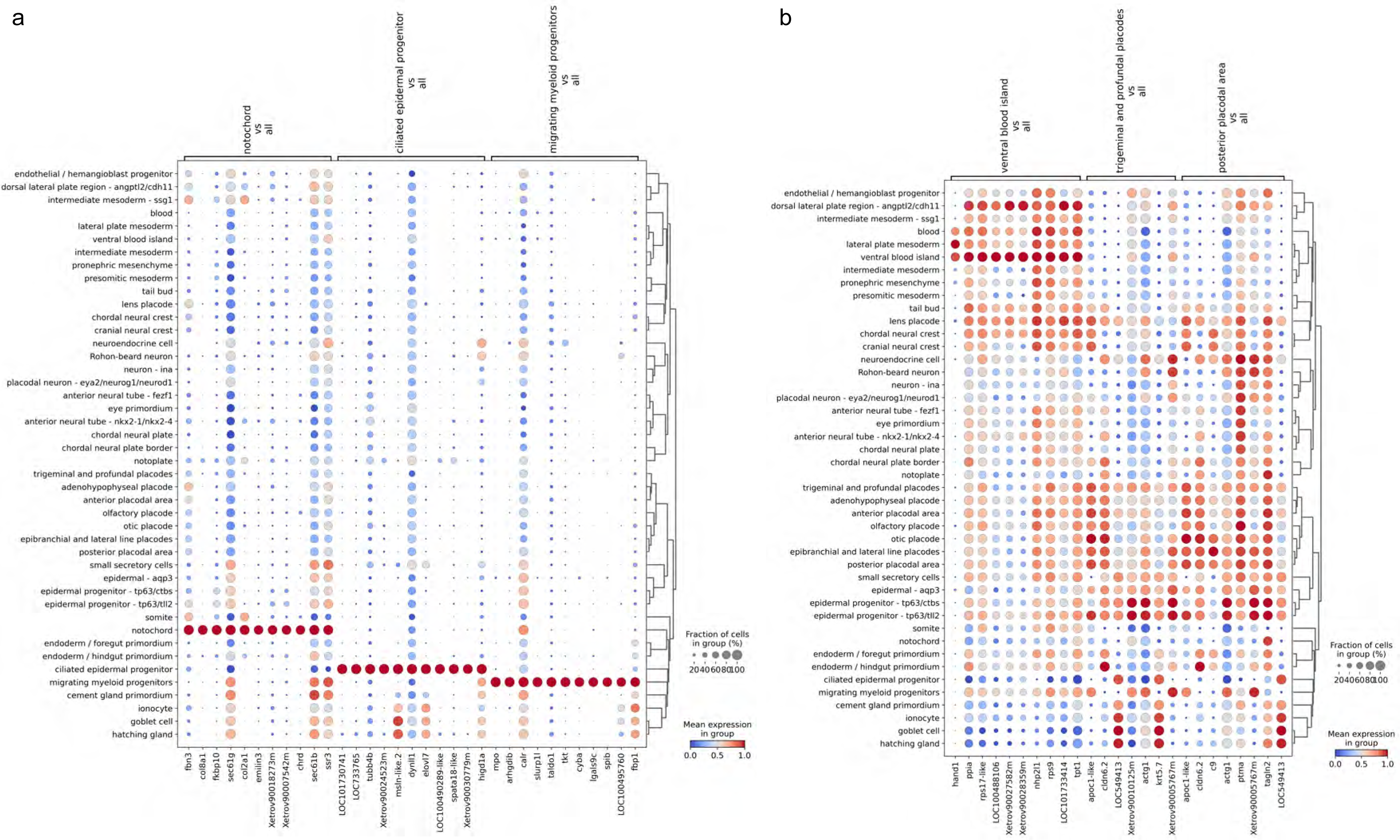
Dotplot of normalized counts for stage NF18. **(a)** The top 10 differentially expressed upregulated genes for the best predictable cell types. **(b)** The top 10 differentially expressed upregulated genes for poorly predictable cell types.

### The amended cell type designation improves the consistency of cell types

Upon further examination of the marker genes, we discarded several cell types which were not self-consistent, i.e. cells and gene markers of this cluster did not help generalize the notion of this type. Specifically we eliminated from further consideration the labels “germ cell” (3 cells total), “trigeminal and profundal placodes” (56 cells), “hatching gland” (129 cells), “epidermal - aqp3” (45 cells) and “neuroendocrine cell” (11 cells) (refer to Fig. 3b, Fig. S2a) as not reliably predictable and not supported by data in the previous edition of the atlas. The respective cells were designated as “Unlabeled”. Additionally, we merged “ventral blood island” (68 cells) and “lateral plate mesoderm” (436 cells) into “lateral plate mesoderm” cluster (Fig. S2b). Having cleaned up the cell type attribution, we re-evaluated the self-consistency of the classification, however the resulting weighted average TPR when predicting cell types by mgB approach remained at 0.6±0.2 (Fig. S4). The performance did not substantially improve since removed cell types had minor contributions to the overall score due to the small respective sub-population size.

### The original cell type designation is robust towards the transition of the genome version

Because the new dataset contains mainly the same cells as the old one we could transfer cell type annotation for persistent cells by matching across the two datasets by cell barcodes. Comparison of cell type predictability across genome versions shows whether gene expression underpinning has changed dramatically, i.e. whether cell types were invariant to the genome annotation and whether new cell types emerged. Focusing on cells which persisted across genome versions, we first evaluated the kNN and the mgB prediction accuracy of cell types in the XA23 the same way we did for the XA18 dataset (Fig. S5). Since some of the marker genes got lost or respective gene sequences and therefore mapped reads changed, the self-consistency of cell type clusters might have suffered substantially. The average cell type prediction TPR for XA23 was 0.8±0.1 by kNN and 0.6±0.2 by mgB. Fittingly, cell type prediction accuracy between XA23 and XA18 didn’t change a lot, TPRs are very similar for both kNN and mgB algorithms (Fig. 4) which confirms that with a few exceptions we kept the key cell type marker genes and cell type clusters.

**Figure 4.**
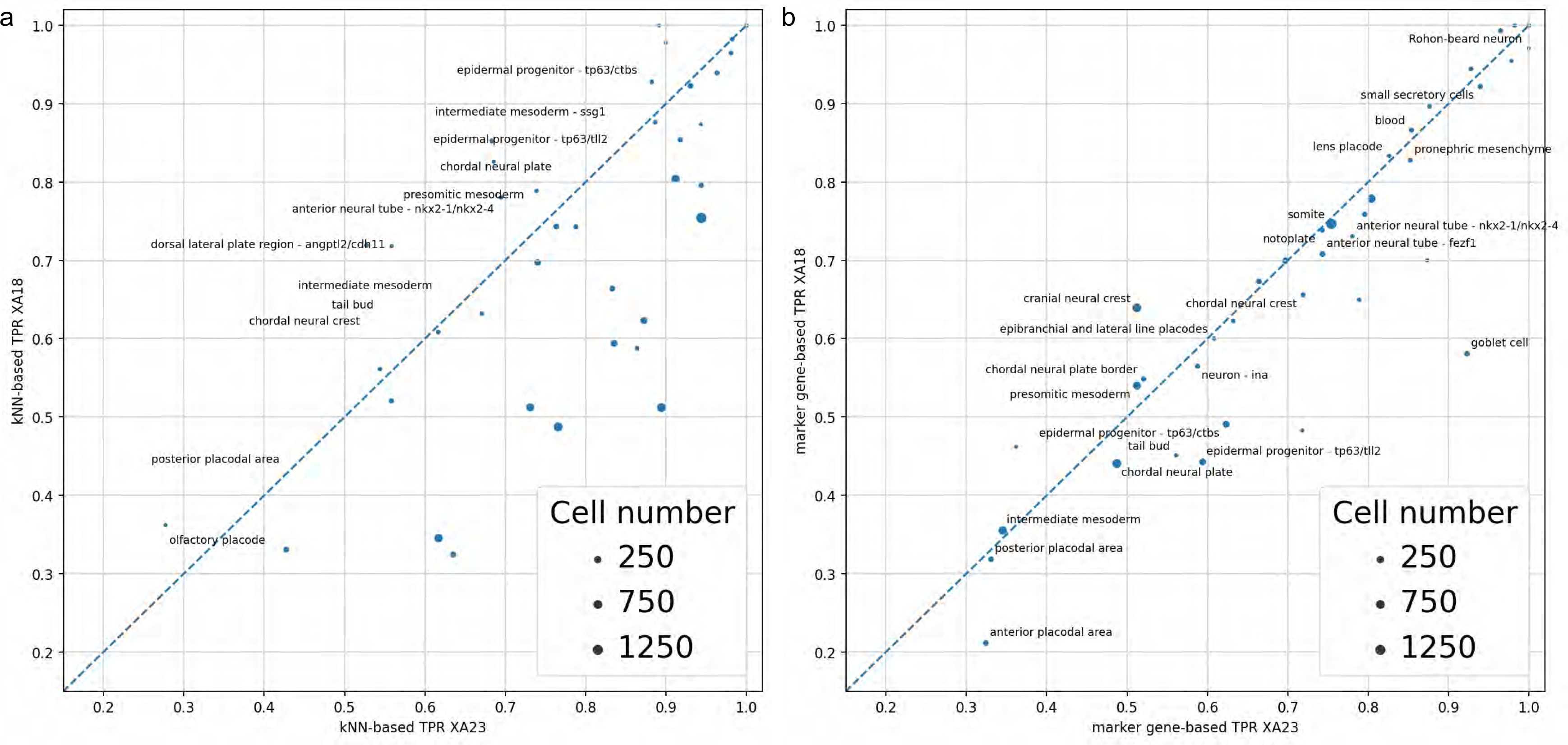
The concordance of cell type predictions across genome versioning: XA18 vs. XA23 cell type prediction TPR: kNN **(a)**, marker gene **(b)**. (Spearman corr. coef. 0.98 for kNN and 0.94 for mgB (both p-values << 1e-10))

Conversely, when comparing differentially expressed upregulated genes in cell types between XA18 and XA23, some new strong (average log_2_FC > 4, fraction of cells expressing > 0.8) marker genes for cell types were found in XA23 in comparison to XA18, while some other were lost (see overall counts in Fig. 5a). This inconsistency is mostly attributed to the discordance in gene models in general and 3’ UTR part of the model in particular since the reads we deal with in this project are primarily 3’ UTR originated. For example, consider two of the genes among the maker genes of “migrating myeloid progenitors“ (Fig. 5b, 5 c): *slurp1l* and *mfsd10.* While *slurp1l is* highly expressed in that cell type when analyzed using genome v9.0, it does not seem to be expressed at all under the annotation v10.7, despite the gene still being present in the genome annotation. The reason for that is the change in 3’-UTR gene annotation (Fig. S6b). Reversely, the *mfsd10*-like gene, an ortholog of MFSD10, got fixed in the new annotation (Fig. S6a), and is uniquely expressed in “migrating myeloid progenitors”.

**Figure 5.**
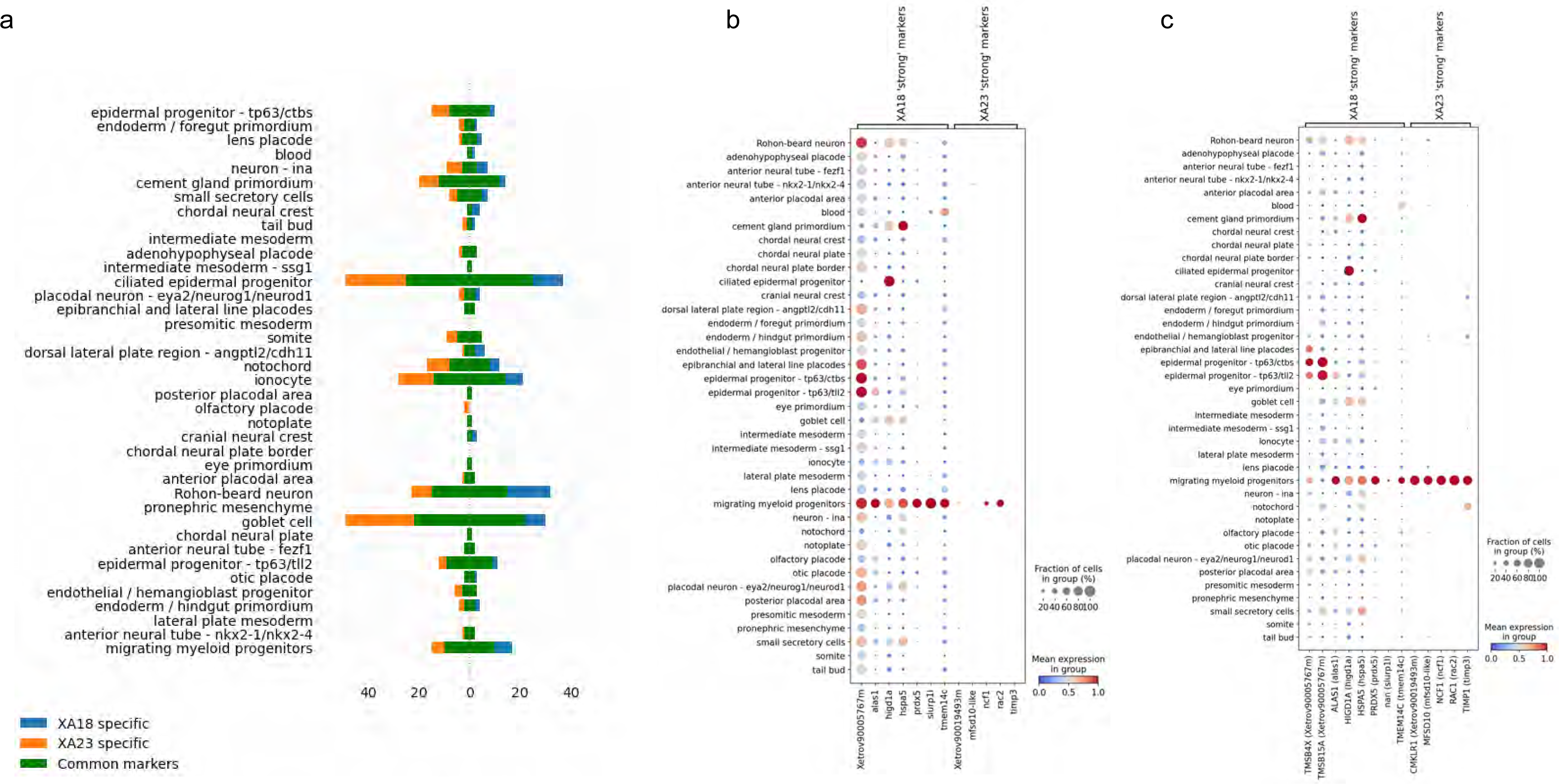
The number of upregulated differentially expressed genes in different NF18 cell types: common between XA18 and XA23 versions and specific for each **(a)**, dot plots of marker genes of “migrating myeloid progenitors” cell type, specific for XA18 and XA23, expression levels in XA18 **(b)** and XA23 **(c)** datasets.

### The new cells provide substantial additional coverage, as exemplified by germ cells

Using our mgB approach, we assigned cell type to the new cells (not presented in the previous version of the atlas) and cells previously annotated as “Outliers”. After re-annotating the new dataset, we looked for clusters with poorly supported annotation, where either our algorithm assigned cells in the same cluster to multiple different cell types, or the cluster consisted of only a small portion of the cell type. Specifically, using this approach we were able to fix the annotation of primordial germ cells (PGCs), which were filtered out in the XA18, because of their low UMI count per cell, possibly because of their transcription silence during early embryonic stages [11]. In total, in the new edition of the atlas, we attributed 477 cells as PGCs, which formed separate clusters at stages 8, 10, 12, 14 and 16. As shown in the dot plot (Fig. 6a), newly annotated germ cells prominently display well-established specific marker genes, e.g. *ddx25, dazl, pgat (LOC100498538-like)* and *dnd1* [12,13,31,14]. While the germ cells were present in the last iteration they were a few and not substantially supported by the marker genes.

**Figure 6.**
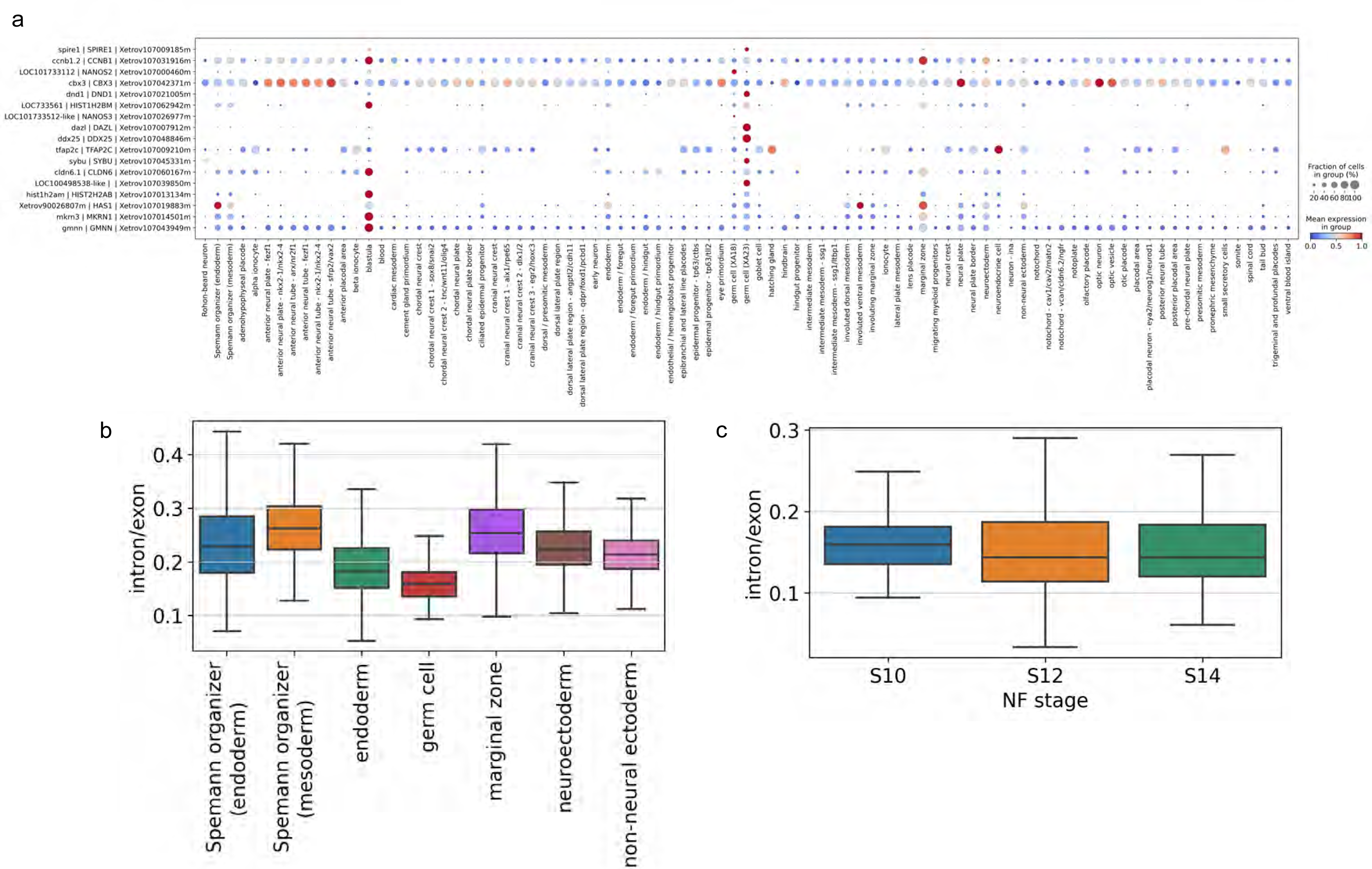
Dotplot of top upregulated differentially expressed genes in germ cells annotated in XA18 and XA23, all stages analyzed jointly **(a)**, boxplot of intron/exon counts ratio for germ cells versus other cell types at the stages observed **(b)** and intron/exon ratio for germ cells at the stages observed separately by inDrops version **(c)**.

Using a substantial population of germ cells, we checked the intron/exon count ratio, that is the ratio of reads which map to introns vs exons within the gene models. With the caveat that gene models are still imperfect, and the read composition is biased towards the 3’ end of the genes, the intron/exon ratio should reflect the level of novel transcription as compared to the presence of maternally deposited, and thus spliced transcripts [15]. Interestingly, we observed that germ cells have significantly lower intron/exon ratio in comparison to all other cell types (Fig. 6b). This difference is very pronounced at the early point St 8 and disappears towards the latest stage where we still have enough germ cells to compute it - St16. This is strong albeit indirect evidence of transcriptional silence since maternal RNA is known to be enriched in spliced RNA [16]. It’s hard to extend the analysis past St18 as the germ cels constitute small proportion of the total population and we simply likely not capturing any.

Note that although the intron/exon ratio in the annotated PGCs changed significantly during the stages observed (the Kruskal-Wallis H-test, see Fig. 6c), it could be affected by the presence of ambient RNA and incorrect gene annotation. Also, the ratio depends on the gene-specific intron capture rate, which depends on the presence of intronic polyA sequences in the gene [17], since the inDrops single-cell technology is based on oligo(dT) RNA capture. Differential expression analysis of annotated germ cells across stages did not reveal any genes to be significantly upregulated with stages, which is in concordance with previously published results [11], where it is suggested that zygotic transcription does not start in PGCs until S14, but contradicts other literature [32], where it is suggested that transcription in PGCs could start as early as at stage S11. On the contrary, several genes which were enriched in the annotated PGCs (e.g. *otx1, ccna1, vegt, velo1, spire1, btg4*) were revealed to be degraded during stages 8-16 (Fig. S16), but this result could also be affected by a high level of ambient mRNA in the assay background.

## Methods

### Embryonic material and sequencing

No new material was collected for this work. Instead, we re-sequenced the RNA libraries for the developmental stages NF11 to NF22 used in (Briggs et al., 2018). Please refer to the methods in (Briggs et al., 2018) for the description of the staging and collection of the embryos, dissociation, cell collection and barcoding (inDrop v2 and v3) and library preparation methods. For re-sequencing, we used two flow cells (two lanes each) of NovaSeq S2 at 100 cycles setting generating a total of 12B reads.

### The reference genome and Gene symbol assignments

In bioinformatics analysis we use the *X.tropicalis* v.10 genome assembly, gene models v. 10.7, specifically we use the file Xentr10-Gene-Sym-HUMAN-BLOSUM45.txt downloaded from Xenbase on Nov 5th, 2021. Thousands of genes in that transcriptome lack interpretable gene symbols, though in many cases an unambiguous protein identity can be identified by sequence homology. To fill the holes in gene symbol assignments we assigned protein gene symbols to each transcript using a modified reciprocal best HMMER3 hit approach [6] based on a target reference set of curated human proteins.

### Single-cell RNAseq analysis

We used cutadapt v.4.4 [19] for splitting libraries by sample barcodes, aligned reads to X. tropicalis v10 genome assembly, gene models v10.7 with STAR aligner v2.710b [20] and estimated intron, spanning and exon UMI counts with DropEst v0.8.6 [21].

For count matrix processing we first corrected raw count matrices for ambient RNA with SoupX v1.6.2 [22], then filtered out cells with less than 150 UMI per cell and 100 genes per cell, and genes being expressed in less than 3 cells and ribosomal genes. Then we used Pearson residuals normalization [23], as implemented in Scanpy v1.9.3 [24] and PCA transformation of normalized counts. Dendrograms for visualization purposes for cell types were built in PCA embeddings using the complete linkage method. For the annotation fullness assessment neighborhood graphs [25] with Euclidean distance on 30 PCs with 15 neighbors were built and Leiden clustering [26] was applied. t-SNE [27] and UMAP embeddings [25] were built for visualization purposes.

Intron/exon ratios were counted across all genes as a ratio of a sum of intron and spanning counts to exon counts for each cell. The Kruskal-Wallis H-test as implemented in *scipy.stats.kruskal* in SciPy v1.10 [30] was used to test if there is a significant difference between intron/exon ratios in germ cells across stages, separately for v2 and v3 libraries.

For pseudo-bulk differential expression analysis of germ cells’ gene expression across stages DeSeq2 normalization and the Likelihood Ratio Test for testing for time-dependence were used [18].

### Cell type annotation assessment with k-Nearest Neighbours Classifier

We used the scikit-learn v1.2 [28] implementation of the kNN classification algorithm (*sklearn.neighbors.KNeighborsClassifier*) with cosine distance in the space of 50 principal components (PCs were obtained as described in the above paragraph), and different k - number of neighbors - parameter values (3, 5, 7, 15) to run 4-fold cross-validation with folds stratified by cell types (*sklearn.model_selection.StratifiedKFold*). For the comparison with the mgB annotation approach, we used kNN results with k=3, since the results didn’t vary as a function of k (data not shown).

### Marker genes selection

For dot plots and marker gene-based cell type annotation we used *scanpy.tl.rank_genes_groups* on log1pCP10k normalized counts to choose marker genes as 10 top-upregulated differentially expressed in cell types’ cells vs. rest cells with Wilcoxon test for p-value estimation and Benjamini-Hochberg correction for multiple testing. All the dot plots were built using *scanpy.pl.dotplot* function on standard scaled by genes log1pCP10k (scaled to a total sum of 10K per cell and log(1+p)) normalized counts.

### Automated cell type annotation

We performed cell type annotation of the new data (“Outliers” cell types in Briggs et al. and new cells from resequencing) using marker genes of cell types annotated in Briggs et al. in the following way. For each cell type, we chose marker genes as the top N differentially expressed genes, filtered by an adjusted p-value and log2foldchange thresholds. After that for the new dataset, we applied over-representation analysis, as implemented in decoupleR v1.4 [29], resulting in cell type activities for each cell, which are equal to -log10(p-value), where the p-value is counted via Fischer’s exact test for the power of intersection between N top expressed genes in the cell and the cell type’s marker genes (we used N=30). Then to get a cell type for each cell we chose the cell type with the highest activity. We verified the approach on the freely available 10x PBMC 3K dataset (*scanpy.datasets.pbmc3k_processed*). The average f1-score weighted by cell-type abundances, with 3-fold cell-type stratified cross-validation was 0.85 (SD 0.08) (Fig. S7). To assign a cell type for each cluster we ranked cell types by their activities in the cluster’s cells and chose the cell type with maximum rank. For the Spearman correlation coefficient, we used *scipy.stats.spearmanr* from SciPy v1.10 [30] with a two-sided alternative hypothesis for p-value calculation.

## Discussion

We presented an updated transcriptomic atlas of developing *Xenopus* embryo at the single-cell level. A number of recent developments enabled a substantial improvement of the atlas in resolution and coverage. First, the new genome release provided for chromosome-level assembly and gene models are much more accurate compared to the previous one. Second, we took advantage of a new batch of data owning to additionally sequenced libraries. Third, the computational pipelines have improved recently allowing to better handle multi-mapped reads [20] and UMI counting [21].

The cell type assignment no longer relies on the error-prone manual process of examining the gene markers. Rather, it is done systematically in a principled way using Machine Learning models. Finally, since the initial release of the embryonic atlas in 2017, the *Xenopus* developmental embryology community has been extensively using the resource while providing us with rich detailed feedback which was taken into account at re-analysis.

We characterized the quality of the atlas in general and specific cell types in particular, pruned some unwarranted cell type labels and rehabilitated others, enriched the representation and augmented the interactive browser with data tracks not provided in the original release. All in all, the new atlas constitutes a substantial step forward and this paper is meant to serve as a companion publication to the atlas itself as well as a re-investigation of the early embryonic differentiation covered by it.

While there is no doubt that this cell atlas will be superseded over the next few years, it will serve as a foundation for designing the forthcoming efforts in single-cell/nucleus profiling, selection of the marker genes for spatial transcriptomics and functional characterization of developmentally important genes and cell types. This atlas will also be federated by cross-specie single-cell resources to enable the comparative perspective onto the evolution of cell types.

## Acknowledgements

This work was supported by NIH OD award R24 OD031956.

We thank Jerome Jullien and Dawn Owens for valuable discussions of our germ cell results.

## Availability of the Data

The Xenopus Embryo Cell Atlas is available at https://tinyurl.com/frogatlas2. The raw and derivative datasets, along with descriptions are under NCBI Gene Expression Omnibus #GSE198491.

The code is available at the GitHub repository https://github.com/Tabula-Rana/XA23.

## Author contributions

Conceptualization: LP

Methodology: KP, MT, AK, LP, AHMB Investigation: KP, MT, LP, AHMB Visualization: KP, LP

Supervision: LP, AHMB Writing original draft: KP, LP

Writing review & editing: KP, AK, LP, AH

## Supplementary Figures

**Figure S1.**
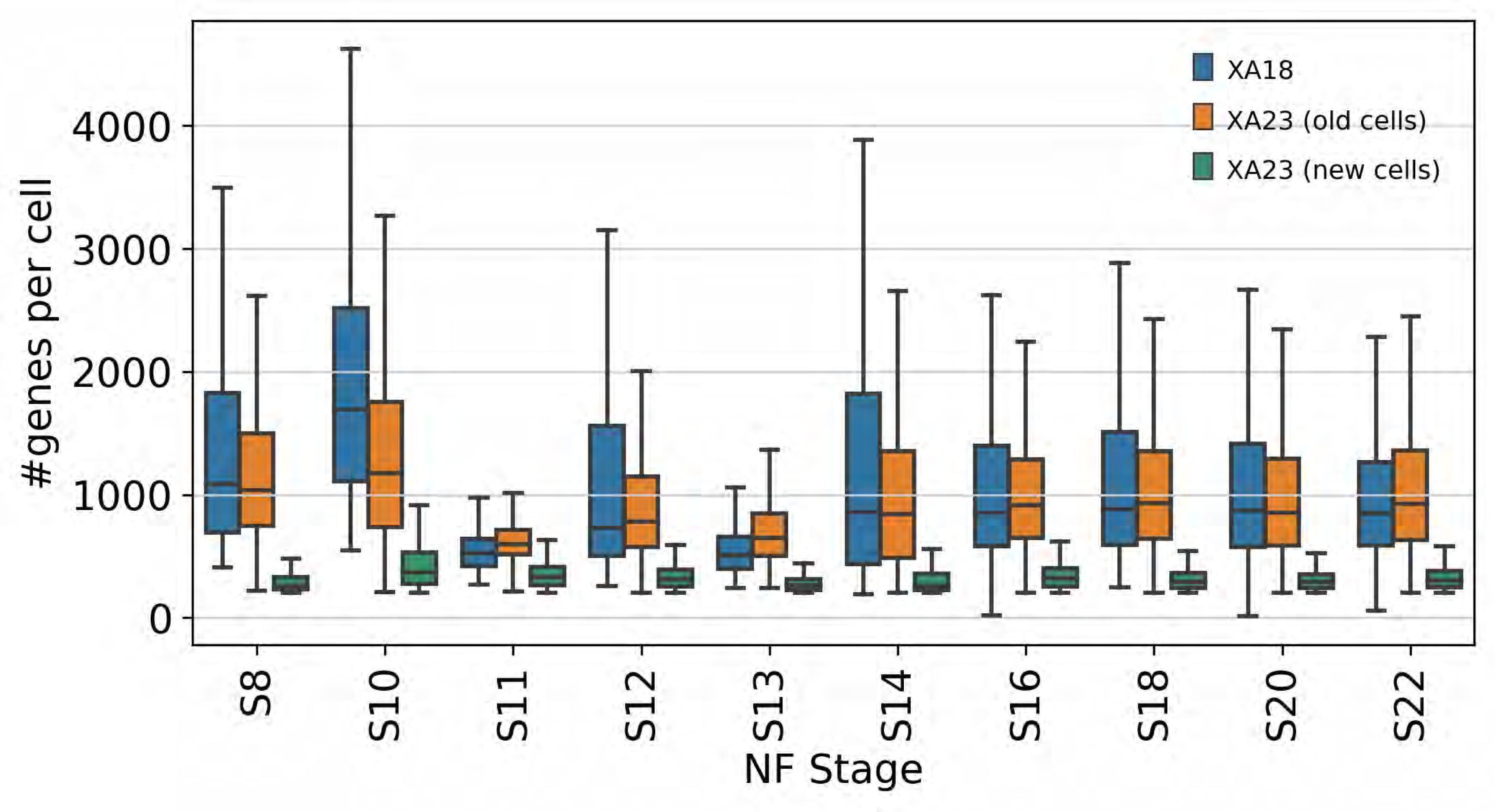
Summary statistics for the number of genes per cell per stage for the XA18 and XA23 datasets (old and new cells separately).

**Figure S2.**
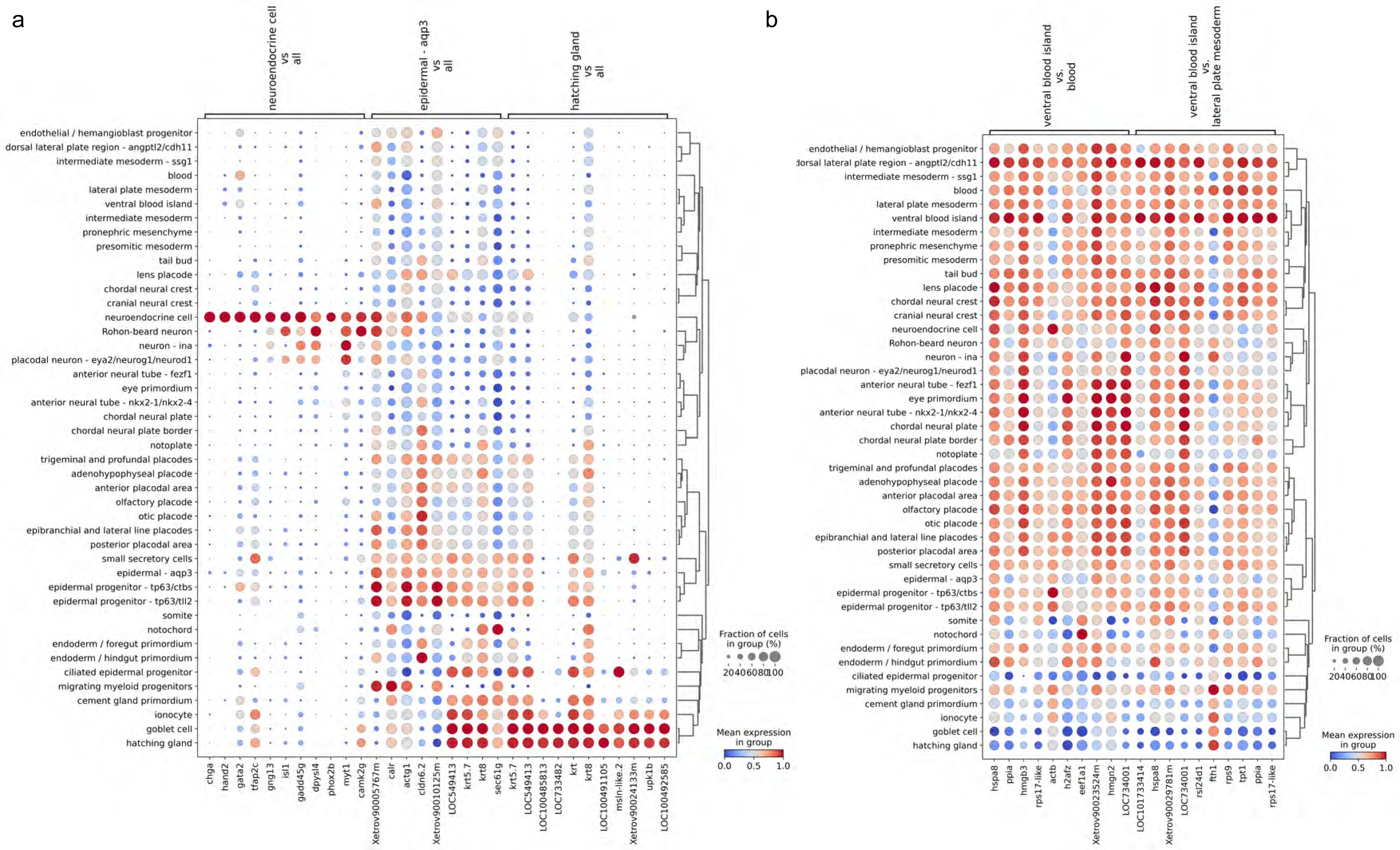
Dot plots for top 10 upregulated differentially expressed genes in cell types poorly predictable with mgB approach **(a)** and top 10 upregulated differentially expressed genes in “ventral blood island” against “blood” and “lateral plate mesoderm” cell types **(b)**.

**Figure S3.**
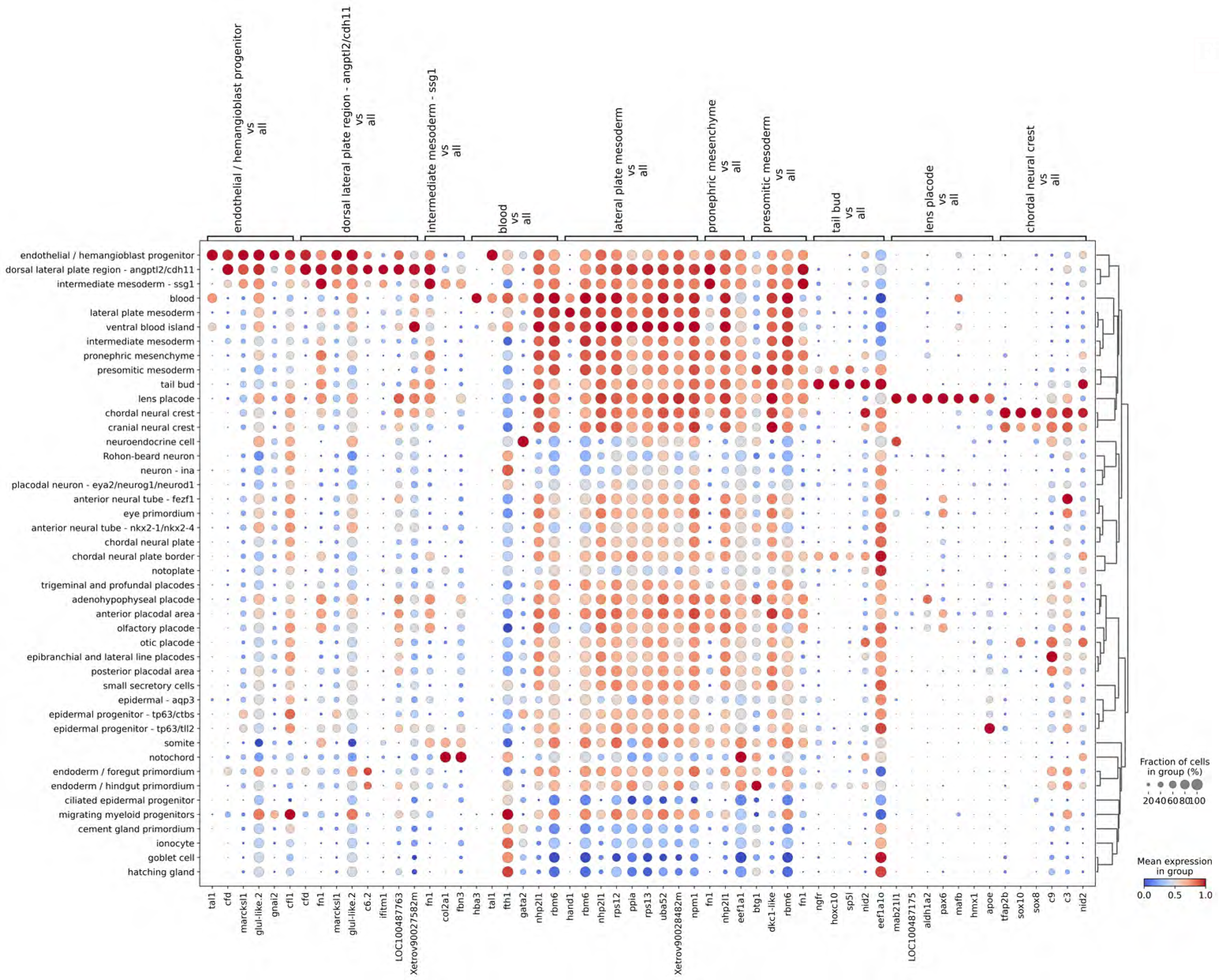
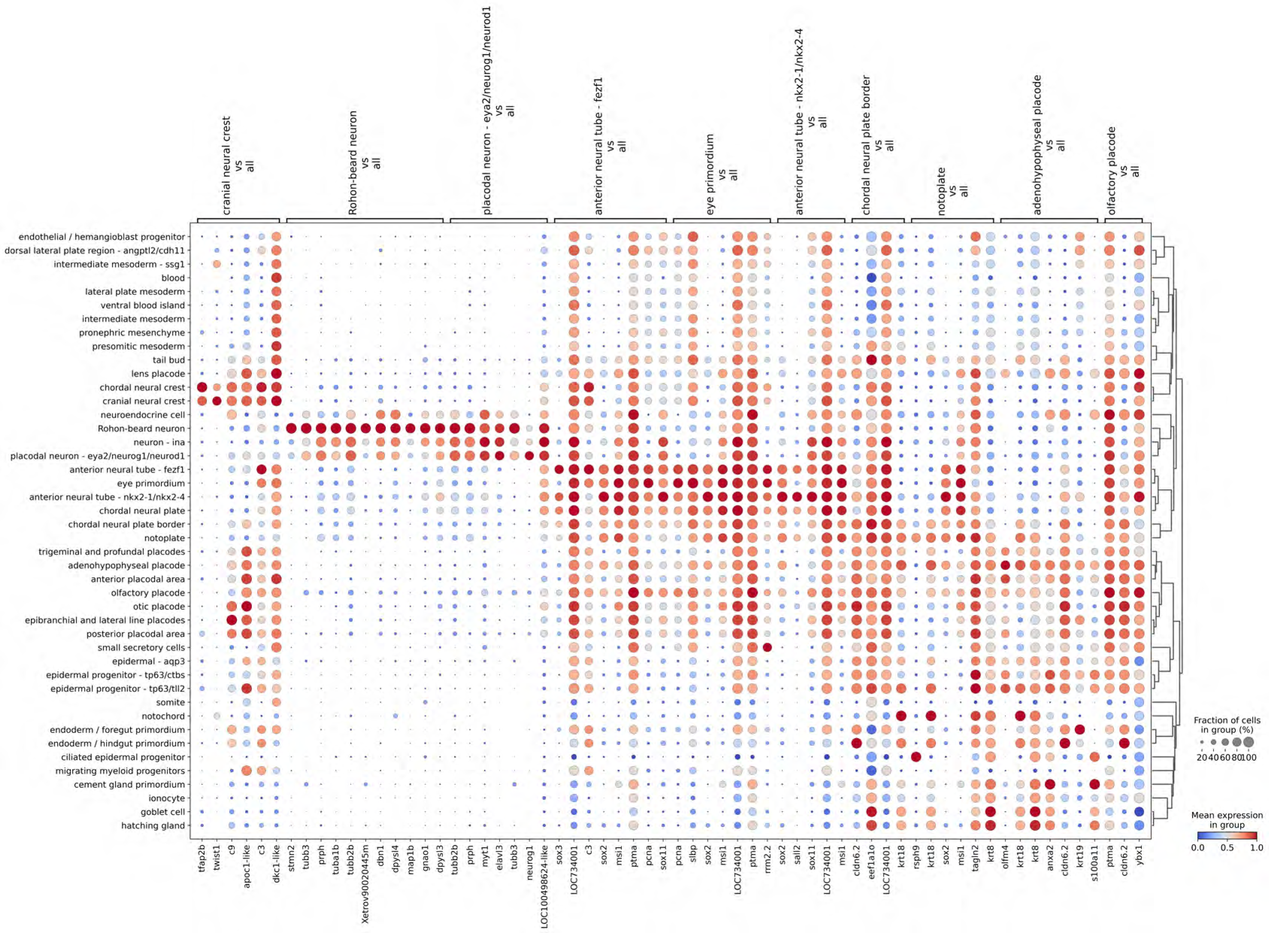
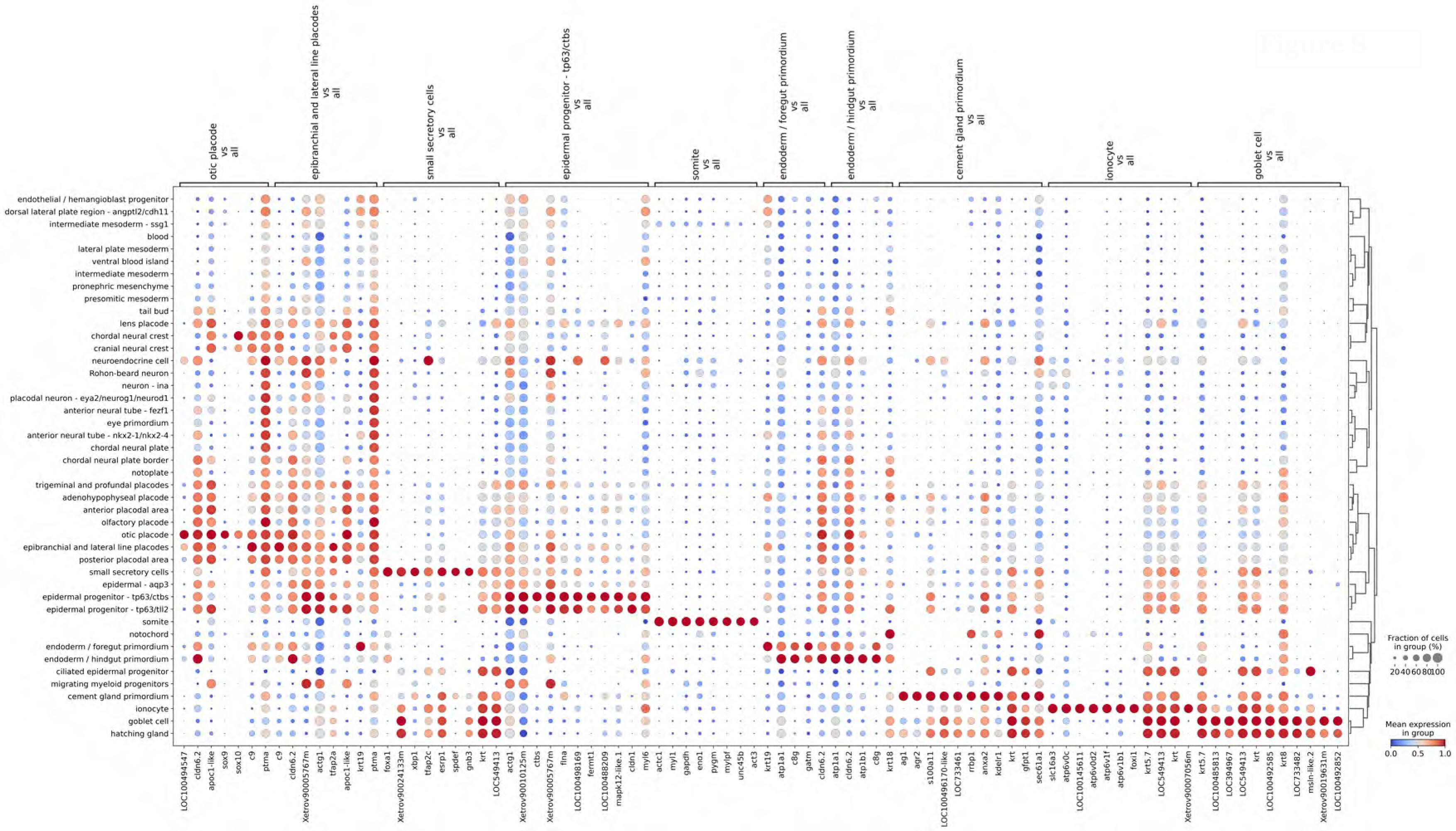
Dot plots of normalized counts of top 10 differentially expressed upregulated genes for “well predictable” cell types.

**Figure S4.**
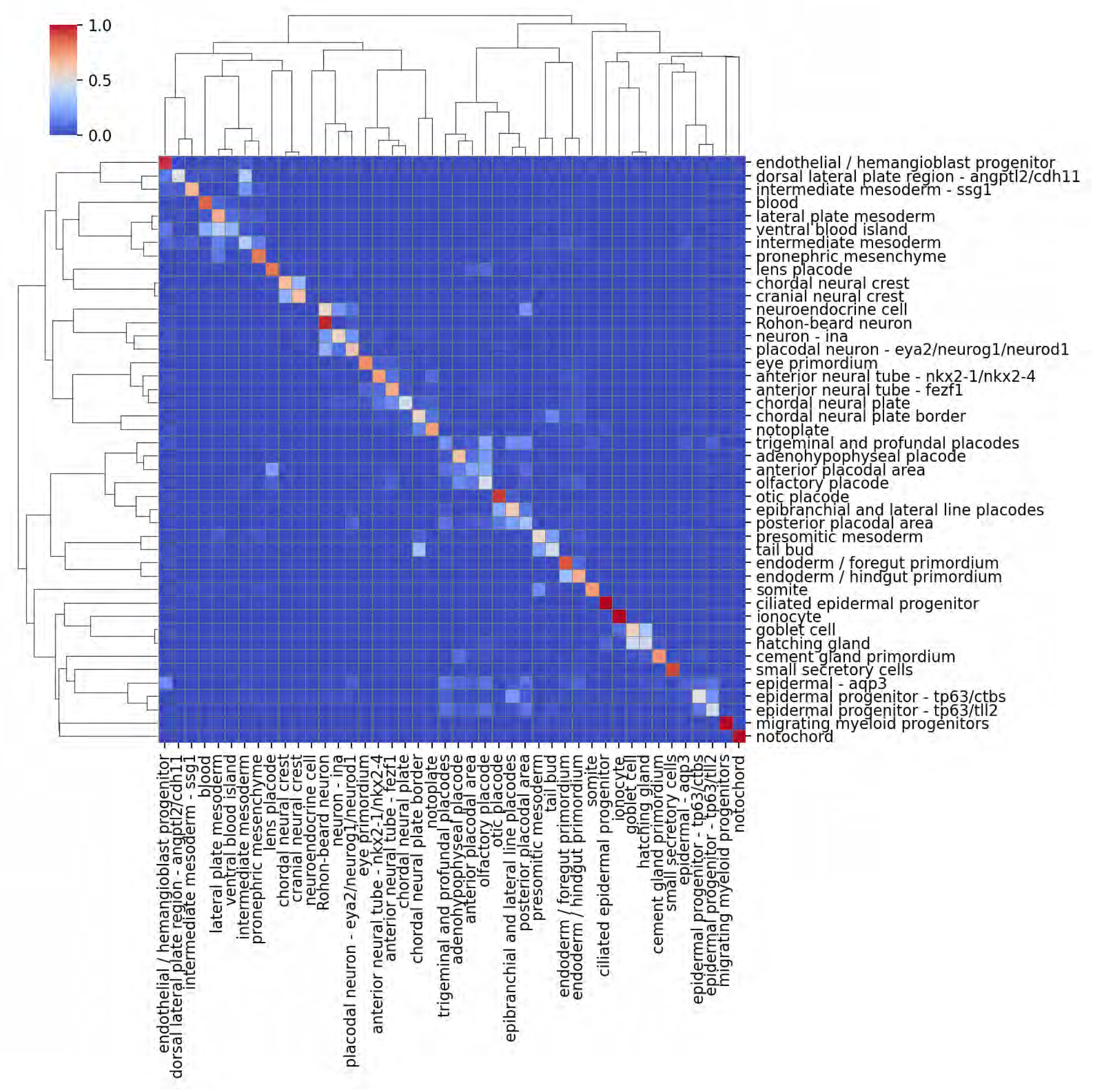
A confusion matrix for NF18 cell type prediction with mgB after discarding poorly predictable cell types.

**Figure S5.**
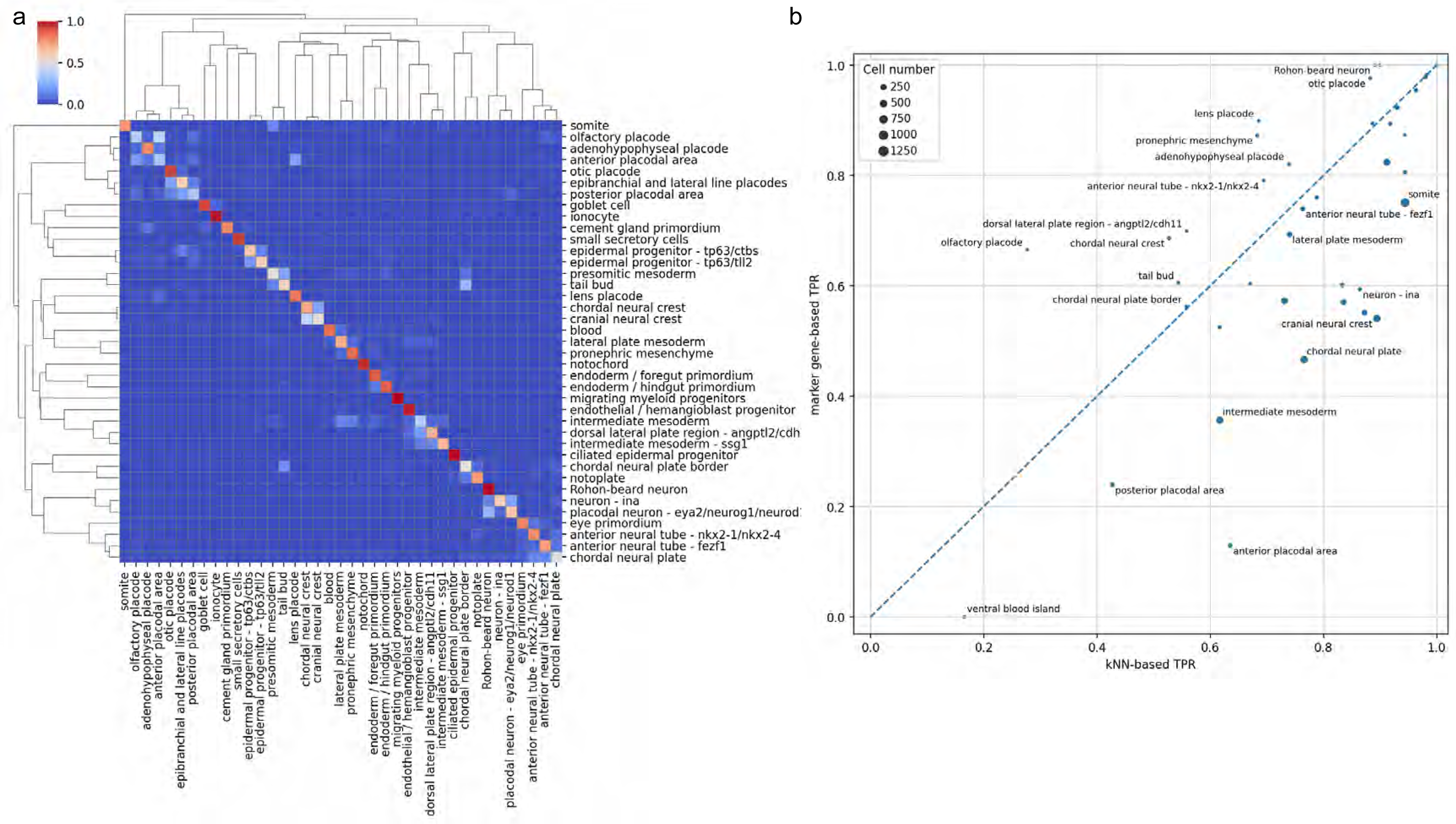
Original cell types prediction accuracy in the new atlas, TPR for mgB vs. TPR for kNN (**a**) and mgB confusion matrix **(b)**.

**Figure S6.**
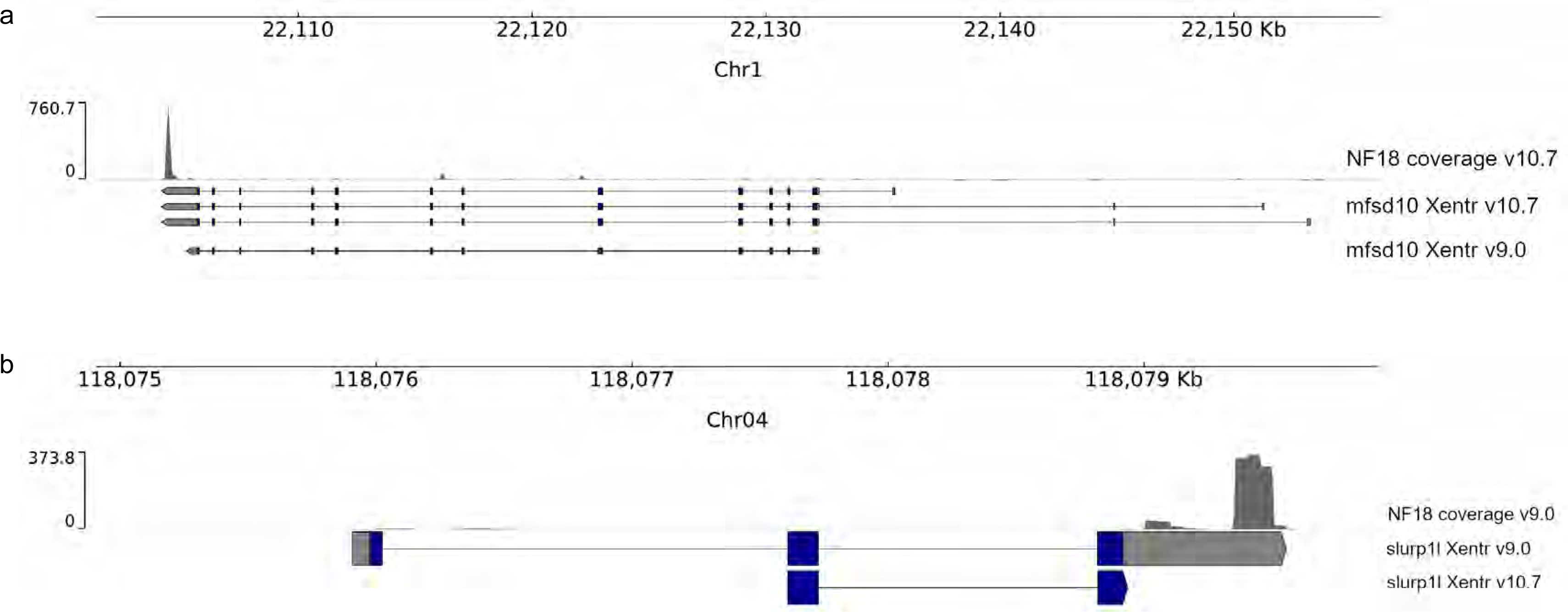
Re-annotation of gene models across genome releases leads to variable results. **(a)** Bam coverage of one of NF18 samples aligned to transcriptome annotation v10.7 (“NF18 coverage v10.7”), *mfsd10* gene coordinates in genome annotation v10.7 (“mfsd10 Xentr v10.7”) and liftover to v10.7 of *mfsd10* gene coordinates in genome annotation v9.0 (“mfsd10 Xentr v9.0”). **(b)** Bam coverage of one of NF18 samples aligned to transcriptome annotation v9.0 (“NF18 coverage v9.0”), *slurp1l* gene coordinates in genome annotation v9.0 (“slurp1l Xentr v9.0”) and liftover to v9.0 of *slurp1l* gene coordinates in genome annotation v10.7 (“slurp1l Xentr v10.7”).

**Figure S7.**
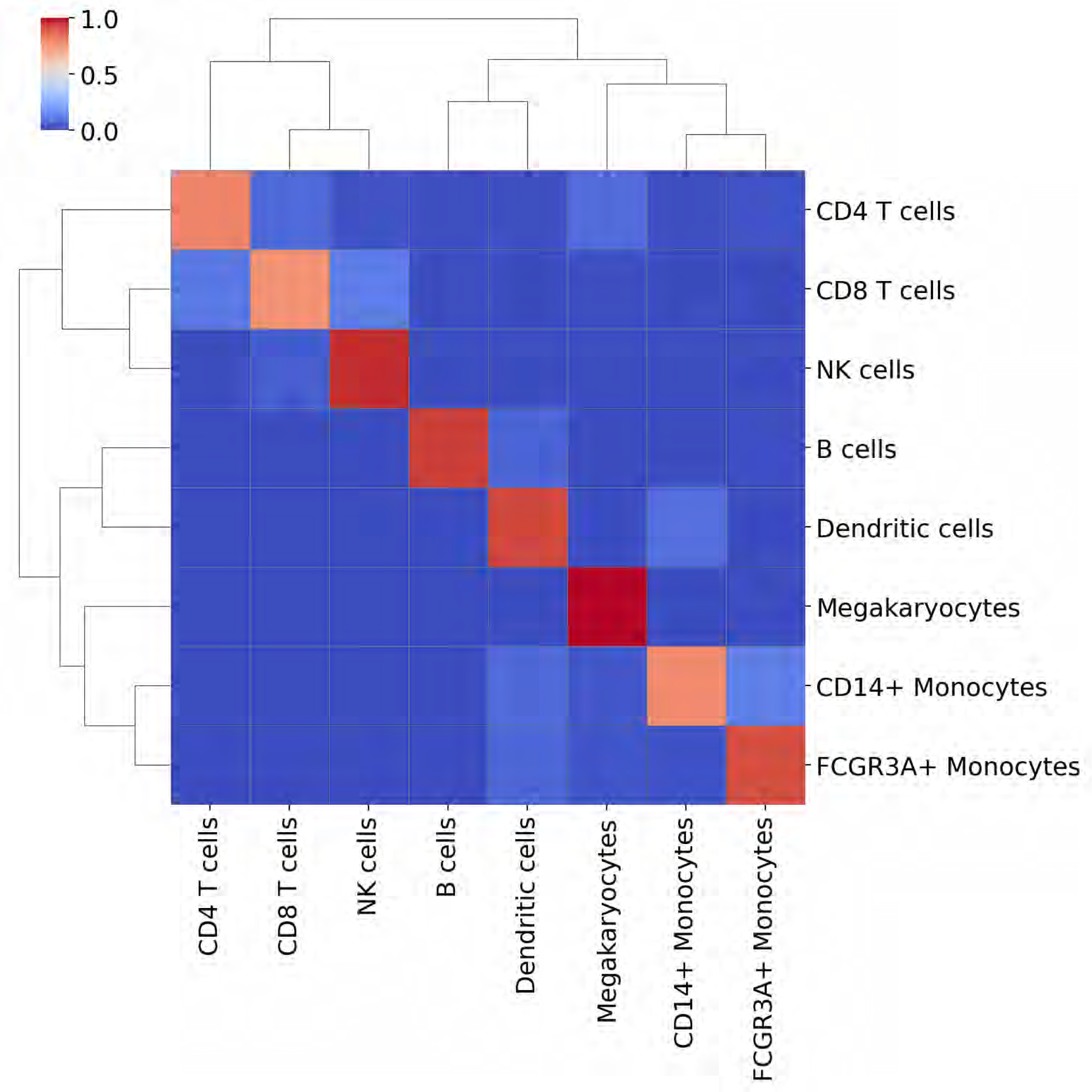
A confusion matrix for the 10x PBMC 3K dataset cell-type prediction via marker gene overrepresentation analysis.

Preliminary PDF with figures is [here](https://drive.google.com/file/d/12tGMl1opMio3pt63YRCMTdsiGHmVcLql/view?usp=drive_link).

